# VTA projections to M1 are essential for reorganization of layer 2-3 network dynamics underlying motor learning

**DOI:** 10.1101/2023.11.22.568212

**Authors:** Amir Ghanayim, Hadas Benisty, Avigail Cohen-Rimon, Sivan Schwartz, Ronen Talmon, Jackie Schiller

**Affiliations:** Department of Neuroscience, Technion Medical School, Bat-Galim, Haifa 31096, Israel; Viterbi Faculty of Electrical and Computer Engineering, Technion, Haifa 3200003, Israel

**Keywords:** primary motor cortex, layer 2-3 pyramidal neurons, VTA, motor learning, network plasticity, two-photon calcium imaging, Riemannian geometry, diffusion map

## Abstract

The primary motor cortex (M1) is crucial for motor skill learning. Previous studies demonstrated that skill acquisition requires dopaminergic VTA (ventral-tegmental area) signaling in M1, however little is known regarding the effect of these inputs at the neuronal and network levels. Using dexterity task, calcium imaging, chemogenetic silencing, and geometric data analysis, we demonstrate VTA-dependent reorganization of M1 layer 2-3 during motor learning. While average activity and average functional connectivity of layer 2-3 network remain stable during learning, the activity kinetics, the correlational configuration of functional connectivity, and average connectivity strength of layer 2-3 neurons gradually transform towards an expert configuration. In addition, task success-failure outcome signaling gradually emerges. Silencing VTA dopaminergic inputs to M1 during learning, prevents all these changes. Our findings demonstrate dopaminergic VTA-dependent formation of outcome signaling and new connectivity configuration of the layer 2-3 network, supporting reorganization of the M1 network for storing new motor skills.

## Introduction

Motor learning is an essential process by which the organism acquires skilled movements and the ability to associate between new sensory information and actions, thus adapting to the ever-changing environmental demands of the world^1–4^.

Learning, planning, and execution of movements, are carried out by a highly complex and distributed system, with the primary motor cortex (M1) being one of its main hubs, generating the output cortical motor commands to the downstream brainstem and spinal cord execution centers^5^. Surmounting evidence supports M1 to be crucial for motor skill learning^6–10^. Previous studies have shown that during motor learning M1 undergoes major plasticity changes, manifesting at multiple levels ranging from motor representations, dendritic computations, branch spike properties, and spine turnover and clustering^6,8,11–14^.

The functional role of M1 in motor control and learning is expected to be cell-type dependent^13,15–18^. Here we concentrate on layer 2-3 pyramidal neurons (PNs), which were shown to represent movement, error estimation, and outcome-related activity^8,13,16,19–22^. In addition, layer 2-3 PNs of M1 were shown to undergo plasticity changes during motor learning both at the activity and structural spine levels^16,22–25^.

Central to motor learning is a reward-based learning process^1^, where reward can in principle inform about the consequences of actions and drive the motor learning process via plasticity mechanisms^26^. The neuronal substrate carrying reward in motor learning and adaptation is the dopaminergic projection systems to both the basal ganglia and cortex^27^. The ventral tegmental area (VTA) was shown to be the main source of dopaminergic projections to M1, innervating dendrites of both superficial and deep layers^28–30^ with forelimb regions being preferentially innervated^31^.

Our working hypothesis is that the dopaminergic VTA projections to M1 directly drive synaptic plasticity mechanisms and the ensuing network connectivity changes in M1, which underlies motor learning. Previous studies have demonstrated the crucial role of dopaminergic signals in M1 for motor task acquisition as well as synaptic plasticity. Destroying the direct dopaminergic projections to M1 eliminated acquisition of a new skilled reaching motor behavior^29^. Dopaminergic system was demonstrated to mediate plasticity changes in both the structural and cellular synaptic plasticity levels of M1 including long term potentiation (LTP) of layer 2-3 horizontal connections, immediate early gene expression, and regulation of spine turnover^26,32–34^. In contrast to the effects of Dopamine at the synaptic level, little is known regarding the effects of the dopaminergic inputs to M1 in altering the activity and functional connectivity of the M1 network during motor learning.

Here, we set out to investigate first, if and how functional activity and connectivity within layer 2-3 network of M1 are altered during learning, second, how the VTA projections to M1 participate in motor learning at the behavioral, network activity and functional connectivity levels, and third, whether VTA projections are essential for the development of outcome signaling in M1. Toward this end, we recorded the activity of layer 2-3 PNs in M1 using two-photon calcium imaging with the genetically encoded calcium indicator GCaMP6s^35^ during learning of a head-fixed hand reach for pellet motor task^10,13^ with and without silencing M1 dopaminergic projections from VTA using DREADDs^36^. We developed a novel non-linear analysis approach to explore the correlational structure of the network throughout training. We used Graph Theory and Riemannian geometry^37–43^ which allow us to compare the functional connectivity of the network at different stages of training. We find that motor learning is associated with gradual and monotonic reorganization of functional connectivity of the M1 layer 2-3 network. In contrast to previous reports^22,44^, we find the majority of neuronal activity as well as the population connectivity gradually transforms towards a new “expert” configuration. Blockade of dopaminergic neurotransmission locally in M1 prevented motor learning at the behavioral level, and concomitantly halted plasticity changes at both the individual neuron activity level as well as at the network activity and functional connectivity levels.

## Results

### Dopaminergic VTA projections to M1 are essential for hand reach motor learning

To study the reorganization of network activity and functional connectivity of layer 2-3 PNs and their dependence on dopaminergic VTA activity in M1 during motor learning, we trained mice to perform a head-fixed version of the forelimb grasping task, where mice learn to reach, grab, and eat a food pellet^10,13^ while performing longitudinal two-photon calcium imaging of GCaMP6s^35^ (Figures 1A-C). To explore the effects of the direct dopaminergic VTA projections to M1 on the motor learning progression and the consequent changes in neuronal activity, we divided the animals into two groups: a manipulated group for which we inhibited these projections for three consecutive training sessions after an initial shaping session, and a second group trained with no manipulations, serving as controls. Dopaminergic projections to M1 were inhibited by DREADDs (hM4D), which were expressed in dopaminergic neurons of the VTA in DAT-Ires-Cre mice. Inhibition was performed by locally injecting CNO to M1 via an access port (CNO sessions) (Figures 1A-E).

**Figure 1:**
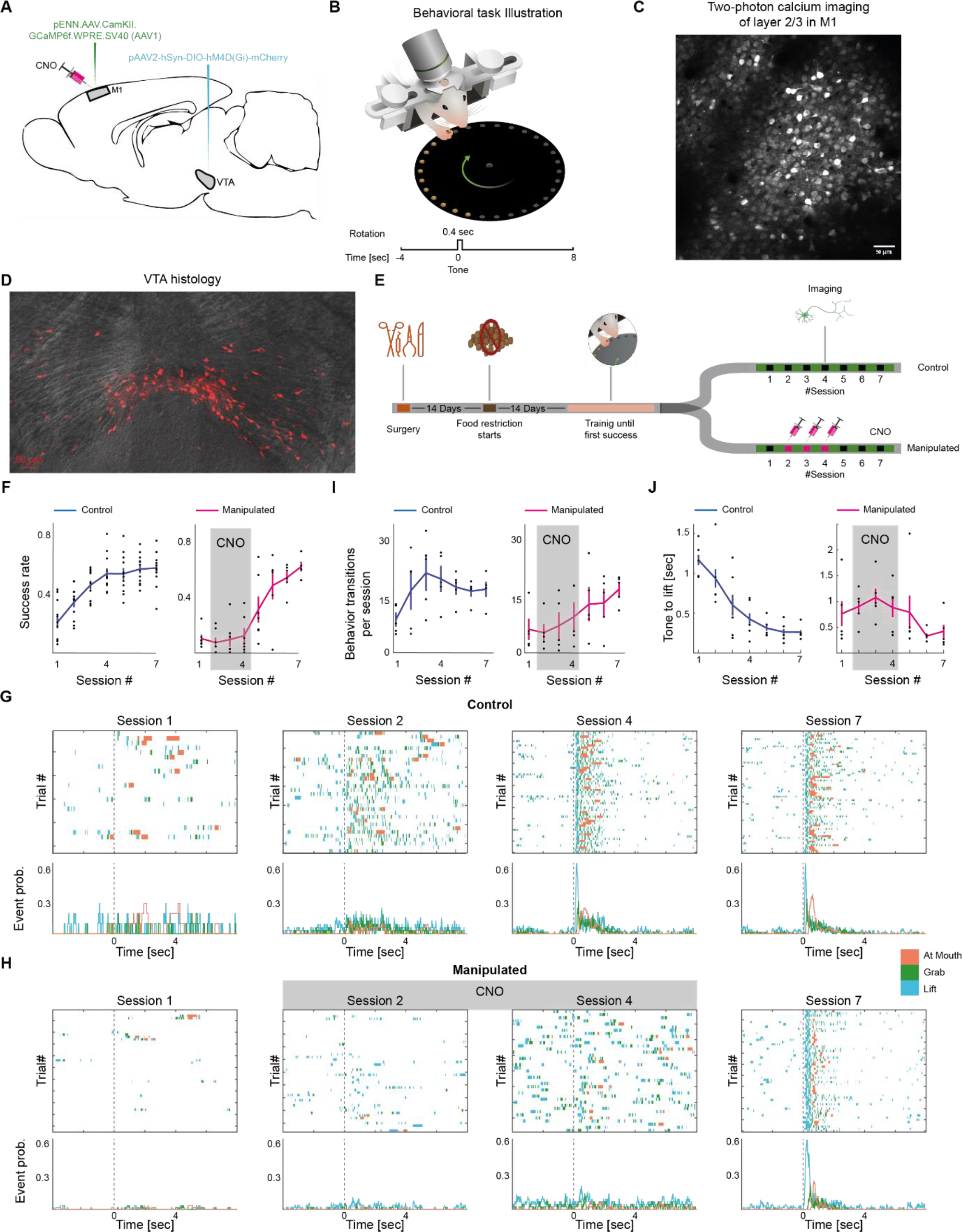
Experimental and behavioral set up for investigating the effect of direct VTA dopaminergic projections to M1 on motor learning. **A**. Schematic of the viral injection of the genetically encoded GCaMP6s to M1 and inhibitory (hM4D) DREADDs to the VTA. CNO was applied locally to M1 via an access port in the relevant experiments. **B**. Behavioral setup. Mice were head-fixed and trained to grab a food pellet positioned on a rotating table. Bottom: trial structure (12 s); tone (go signal) after 4 s, plate rotates and pellet reaches position at tone+0.4 sec. **C**. Example of a typical field of view showing average projections of two-photon imaging of GCaMP6-positive layer 2-3 neurons (250 μm from pia) of an M1 field. **D**. A coronal section of hM4D DREADDs staining with mCherry (red) in the VTA area (red neuronal staining merged with a transmitted light image). **E**. Experimental design for learning a motor task for the control and manipulated mice groups during VTA silencing with local CNO in M1. **F**. Population and average data (mean±SEM) of success rate (fraction of successful trials) per training session during learning, n=12 for controls and n=7 for manipulated mice (p=7.4·10^-7^ unpaired t-test comparing success rate in the 3^rd^ session of manipulated group n=7 mice to control group, n=12). **G**. Example ethograms in control mice annotating the behavior during training over consecutive trials presenting ‘Lift’, ‘Grab’ and ‘At Mouth’ as a function of time on training sessions 1, 2, 4 and 7. Bottom panel, empiric probabilities for events corresponding to the ethograms. See also Figure S1 for additional animals from both groups. **H**. Same as I for a manipulated mice (CNO at sessions 2, 4). **I**. Population and average data (mean±SEM) presenting number of transitions between the five behavioral states (hand lift, reach, grab, supinate, at mouth) divided by the number of trials per training session, n=5 for controls and n=5 for manipulated mice. **J**. Population and average data (mean±SEM) presenting time delay from go cue (tone) to ‘Lift’, n=5 for controls and n=5 for manipulated mice. For Figures 1F, I, and H black dots are population data (per animal), blue and pink are mean (±SEM) across animals, n=12 for controls and n=7 for manipulated mice.

The control animal group demonstrated a steady and gradual improvement in motor execution of the task until reaching a stable level. We quantified task performance by the success rate and by evaluating the sequence of motor events during learning. To calculate the success rate, we determined trials successful if animals succeeded in grabbing and consuming the pellet. We observed a gradual increase in success rates at the first four training sessions (R^2^=0.55, p=2·10^-9^), after which success rate did not change significantly (p=0.691 for one way ANOVA comparing success rate in sessions 5 through 7, n=12 mice) where all animals maintained high proficiency (of 0.54±0.03 at 7^th^ session, n=12 mice, Figure 1F; Videos S1-3). Learning as evaluated by success rate was impaired and remained low for the manipulated group in sessions where CNO was injected (Figure 1F; Videos S4-5; 0.08±0.03, p=7.4·10^-7^ unpaired t-test comparing success rate in the 3^rd^ session of manipulated group n=7 mice to control group, n=12). In consecutive sessions, once CNO was lifted, learning progressed gradually (linear fit for training sessions 4-6, R^2^=0.4, p= 0.0003; success rate at 7^th^ session 0.44±0.05, n=7; Video S6). Overall, the manipulated group reached a slightly lower success rate at the 7^th^ session but there was no significant difference compared to the control group (p=0.11, unpaired t-test). This small difference possibly relates to the smaller number of post CNO training sessions compared to the control mice training sessions.

To control for any direct inhibitory effect of the CNO injection unrelated to its effect via the expression of hM4D DREADDs in the VTA, we injected CNO locally to M1 in mice that were not injected with the hM4D expressing virus to the VTA. Comparing the CNO injected groups with and without the hM4D viral expression reveals significant differences in the CNO injected sessions (Figure S1A). While the CNO injected mice without viral injection progressed in the learning, the manipulated group with viral expression differed significantly and did not show progression in the success rate (p=1.8·10^-4^, unpaired t-test for the three CNO injected training sessions between the viral injected and non-viral injected groups). In addition, we did not see significant differences between the CNO injected mice in the non-viral injected group and the control mice where no CNO or virus was injected (one way ANOVA p=0.18). These results indicate that the effect of CNO injection on motor learning progression is mediated via the hM4D DREADDs expression in axons projecting to M1.

In addition to success rate, we identified the behavioral events during each training session (Lift, Reach, Grab, Supinate and At-Mouth, see Methods) using a semi-supervised software^45^ as exemplified in Figures 1G-H and Figure S1B-C. We next evaluated the number of transitions in behavioral states per trial at each training session (Figure 1I). For the control group we observe a gradual increase in the first three training sessions (R^2^=0.3, p=0.03, n=5 mice for linear fit), after which the number of transitions slightly decreased and then stabilized (no significant change during sessions 4 through 7, p=0.72 for one way ANOVA, n=5 mice). For the manipulated group, the number transitions remain low in the first 3 sessions (6.2±2.7 transitions, p=0.79, tailed t-test comparing the 1st training session to the 3^rd^, n=5 mice) and was significantly different compared to the control group (p= 1.9·10^-4^, unpaired t-test comparing number of transitions in session 2-4 for the manipulated group vs. the control group. In addition to the number of transitions, we evaluated the time delay between the sensory cue (tone) and the first Lift event along the training sessions (Figure 1J). We found that while in control mice this time delay decreases during the first four training sessions and then levels off, for the manipulated group the time to lift remain high during the CNO training sessions, and decreases in training sessions where CNO is lifted (p=0.0086, for unpaired tailed t-test asking if delay time in sessions 2-4 is longer than delay time in the 7^th^ session). These results suggest that direct VTA dopaminergic projections to M1 play a significant role in learning of a motor sequence^46^.

### Re-organization of layer 2-3 neurons’ activity during motor learning and the role of direct M1 dopaminergic projections in the process

We explored changes in the activity of layer 2-3 neurons during the training sessions. When considering the overall mean activity (cells across trials, per training session) we found it did not change significantly with training for both control and manipulated groups^16,24^ (Figure 2A, p=0.97, n=12 control group; p=0.38, n=7 manipulated group, one way ANOVA test).

**Figure 2.**
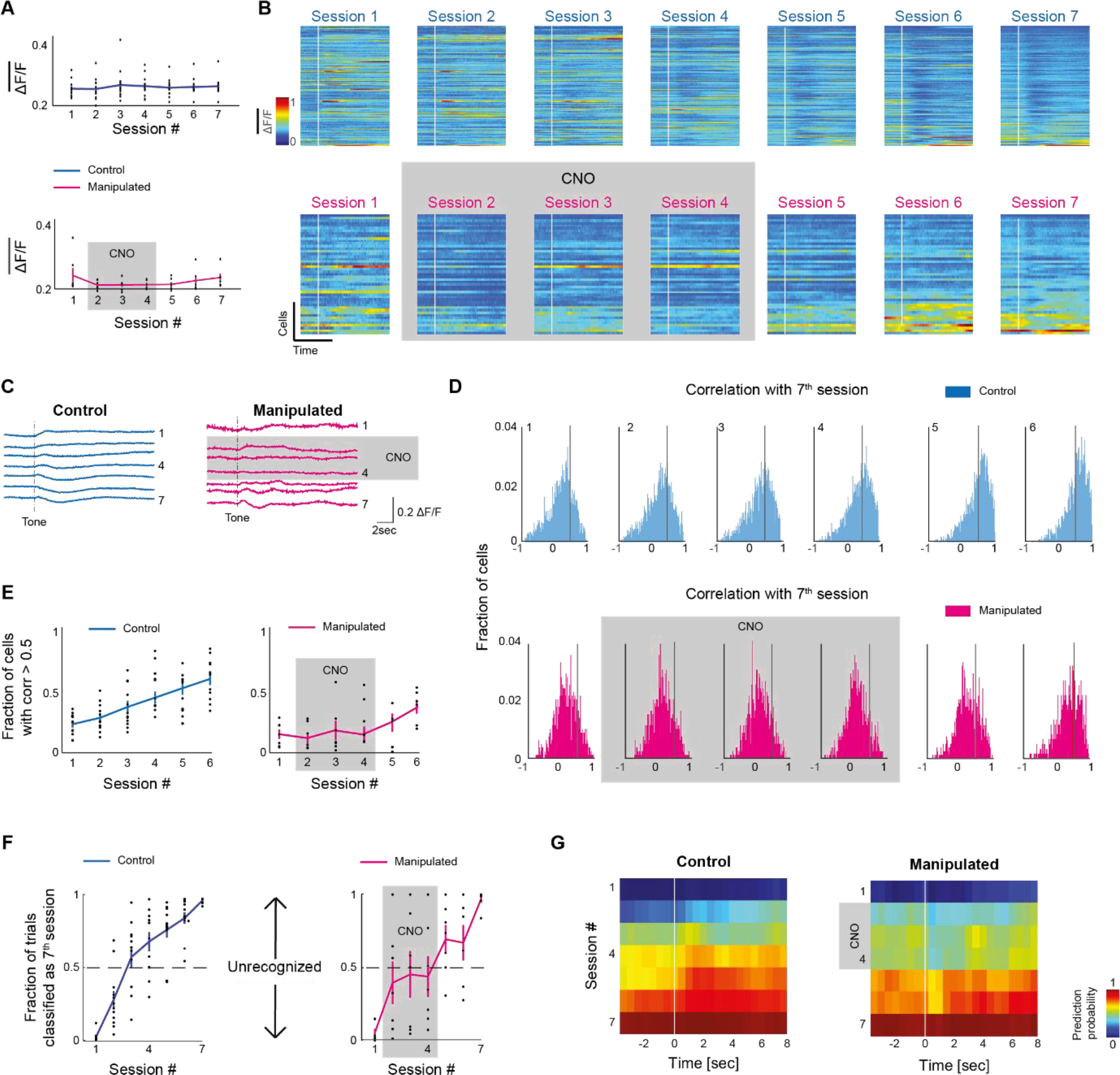
Reorganization of layer 2-3 PN activity in M1 during motor learning at the single neuron and the population levels. **A**. Population data showing average activity (across cells, time, and trials) per training session. **B**. Examples of individual neuron activity vs. time, averaged across consecutive trials per training session (1-7). Color encodes the percent change in fluorescence (ΔF/F). **C**. Example traces of the mean activity along the trial time (average across cells and trials) per training session. Same mice as in B. **D**. Population data showing histograms of correlation values between the average activity trace of each cell per session and its own average trace in session 7. Gray vertical line indicates correlation=0.5. **E**. Fraction of cells having correlation >0.5 with their average activity trace in session 7 (corresponding to data in E). **F**. Fraction of trials being classified as expert by an SVM binary classifier trained to separate ensemble activity of beginner profile from expert profile, trained using activity profile averaged across a 1 sec time window centered at t=1sec. **G**. Heat map showing probability for trials to be classified as expert (vs. beginner) based on ensemble activity for a sliding window analysis (window size 1sec, window hop 0.5sec), n=12 for controls and n=7 for manipulated mice. Results for sessions 1 and 7 are a product of 10-fold cross validation. For days 2-6 results are a product of the trained models. For figures 2A, E, and F, black dots are population data (per mice), blue and pink are mean (±SEM) across mice, n=12 for controls and n=7 for manipulated mice, correspondingly.

While these results might indicate that M1 layer 2-3 network activity remains unchanged with training, a closer look at the dynamics of the activity profile of the cells throughout the training sessions reveals that the network undergoes a gradual reorganization, with an emergence of a tri-phasic average activity pattern^13^. Examples of the mean activity of cells (across trials), per session, sorted by their variance at the 7^th^ session (expert), extracted from a control animal and for a manipulated animal is shown in Figure 2B and their corresponding average activity traces across cells and trials per training sessions (Figure 2C). To quantify this transformation of activity dynamics during training, we calculated the correlation coefficient between the average activity traces of each cell in the different training sessions with its own expert trace of average activity at the 7^th^ session (see Methods). For the control group we observed a shift towards higher values with training, indicating that a growing number of cells adopt a dynamic pattern, similar to their individual dynamics as experts (i.e., 7^th^ session, Figure 2D). For the manipulated group, this process emerged only at the 5^th^ session as training with no CNO is resumed, though still did not reach a comparable value to that of the control (Figure 2D). We quantified this process by counting the fraction of cells with correlation coefficient higher than 0.5 (Figure 2D, grey line) in each training session. We found a gradual increase in the control group (R^2^=0.45, p=1.4^-10^, for linear fit) while for the manipulated group an increase was observed only after lifting the CNO (Figure 2E).

To further quantify the reorganization of the layer 2-3 PN population towards a specific activity profile throughout the training process, we trained a linear classifier to separate between the activity profile of the 1^st^ session (beginner) and the 7^th^ session (expert). The classifier was based on the ensemble activity profile averaged across a short sliding time window throughout the trials (1 second time window, with 0.5 seconds window hop, see Methods). We then applied this classifier to sessions 2-6 and counted the fraction of trials in each session that were classified as expert (Figure 2F for a time window centered at go cue+1sec and Figure 2G for the entire trial time, window length=1sec, hop=0.5 sec, see Methods). For the control group we observed a monotonic increase of trials being classified as expert, indicating a gradual and steady shift of population activity towards an expert profile. Repeating this analysis for the manipulated group revealed that CNO injection leads to an unrecognizable activity profile as trials were not consistently classified as either beginner or expert. Once CNO was lifted at the 5^th^ session, most trials were classified as expert, indicating transformation of the neuronal population activity toward the expert configuration.

Together, these results highlight the crucial role the dopaminergic innervation in M1 plays in the gradual development of the learning related dynamics at the population of layer 2-3 neurons.

### Development of outcome related signaling in layer 2-3 network of M1 during motor learning and the role of direct M1 dopaminergic projections in the process

Our previous study showed a specialized sub-population of layer 2-3 neurons, which reported the outcome of the hand reach trial. These outcome related neurons developed during the learning process^13^.

Here, we studied the involvement of M1 VTA projections in the development of the outcome signal. We calculated the fraction of indicative neurons that reliably report the outcome of the trial^13^ (we defined ‘‘indicative neurons’’ as neurons that predict success or failure trials with 99% confidence) at the different training days in both the control and manipulated groups (Figure 3A). Similar to our previous results^13^ we observed a gradual increase in the fraction of indicative neurons along the training sessions. In contrast, in the manipulated group we did not observe the emergence of indicative neurons during sessions where dopaminergic VTA axons in M1 were silenced (Figure 3A). An increase was observed only after lifting the CNO in subsequent training sessions (Figure 3A).

**Figure 3.**
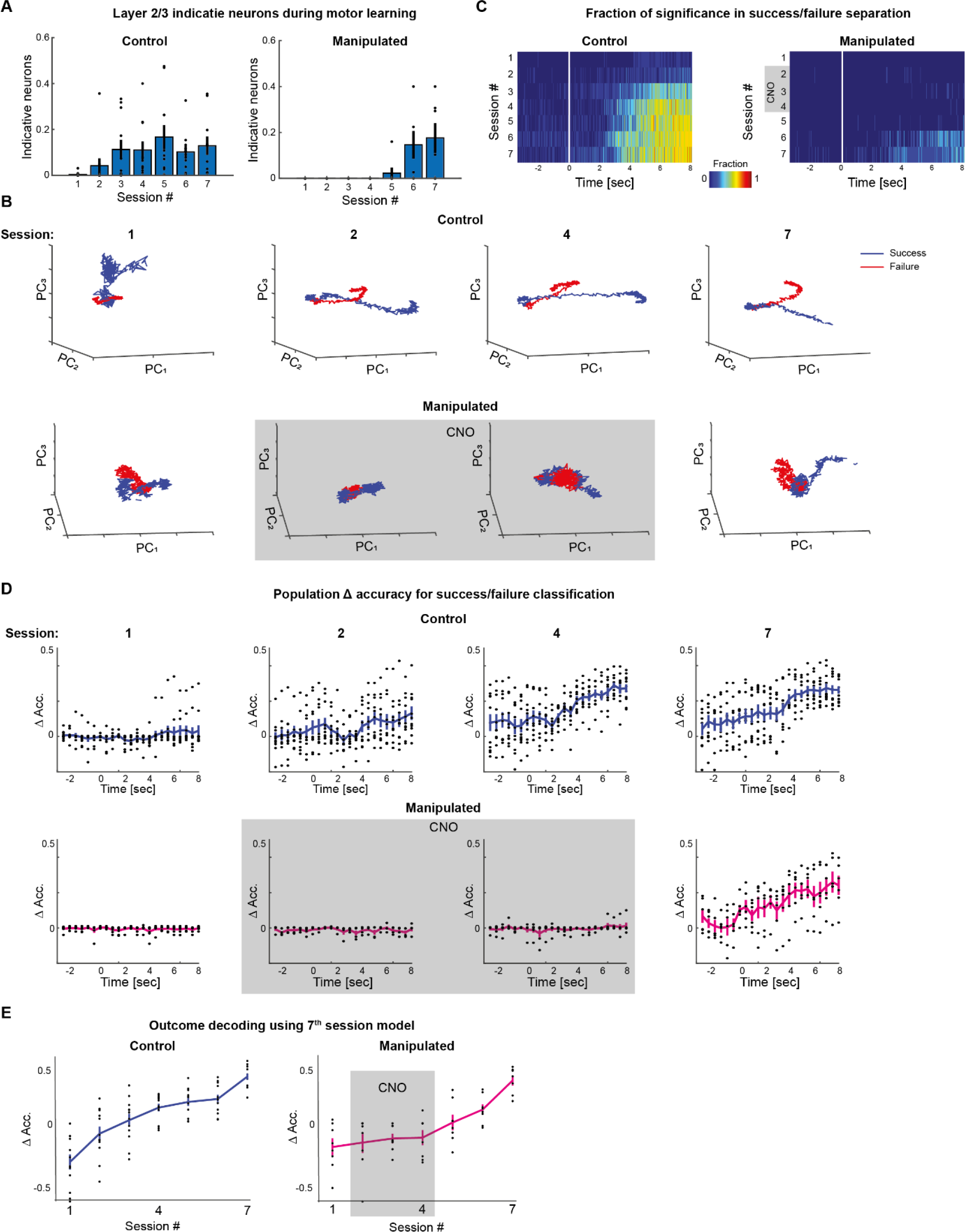
Development of success/failure signal in M1 network during motor learning in control and manipulated mice. **A**. Average fraction of indicative neurons (mean±SEM) per training session (n=12 for controls and n=7 for manipulated mice). **B**. Examples of temporal network dynamics during four representative training sessions (1,2,4,7) as captured by the 3 principal components averaged across trials according to outcome: success (blue) and failure (red). Top panel, control mouse, bottom panel manipulated mouse (grey box indicating CNO sessions). **C**. Fraction of animals where network dynamics showed significant difference between successful and failure trials, as a function of time, per training session (see Methods) for the control group (left) and manipulated group (right). **D**. Population data showing delta accuracy rate (accuracy minus chance level) for classification of outcome (success/failure) based on the ensemble activity of entire population of neurons in a sliding window (window length 1sec, window hop 0.5 sec), shown for four representative sessions (1,2,4,7). Top, control group and bottom, manipulated group. **E**. Delta accuracy for classification of outcome (success/failure) using models trained on the final analysis window (go cue+7 to go cue +8 sec) of expert sessions (7th) and applied to all other sessions. For the control group (left), p=8.5·10^-4^, p=1.0·10^-4^, p=6.3·10^-5^, p=4.0·10^-9^ for sessions 4-7 respectively, tailed t-test, n=12. For the manipulated group, p=0.013 for the 6^th^ session and p=7.4·10^-5^ for the 7^th^ session. Results for 7^th^ session are the product of 10 fold cross-validation. For figures 3D, and E, black dots are population data (per animal), blue and pink are mean (±SEM) across animals, n=12 for controls and n=7 for manipulated animals. See also Figure S2. * Indicates p<0.05.

We next evaluated the outcome signals at the population level using a dynamical system analysis as we described previously^13,47,48^. We extracted principal components of the ensemble activity to capture 95% of variance, and then averaged the PCs across trials according to outcome to obtain two trajectories per session: one representing the average ensemble dynamics in successful trials and the other representing the average dynamics in failed trials (Figure 3B). We quantified the distance between the success and failure trajectories as a function of time and used permutation testing to determine whether these differences are significant (see Methods). We counted the fraction of animals where success-failure trajectory distances were significant (p<0.05, see Methods; Figure 3C). In the first training session the average dynamics did not differentiate by outcome for both control and manipulated animals (Figures 3B-C). With training, the trajectories of the control group became increasingly separated by outcome mostly after the movement time window. Significant differences emerged in sessions 2-3 and onward, at the post movement time segment (5-6 seconds after the go cue) (Figures 3B-C and Figure S2A), which indicates that the population activity adopted a different dynamic for success compared to failure trials. In contrast, for the manipulated group, the separation between success and failure trajectories did not emerge during training sessions with CNO injections (2^nd^ -4^th^ sessions), but rather, it emerged in training session only when CNO is lifted but remained lower than the separation seen in the control group did not reach separation levels seen in the control group.

We further quantified the extent of which outcome can be decoded from the entire population activity of layer 2-3 PNs by training a set of binary classifiers (see Methods). Sliding the analysis window across trial time (1 sec time window and 0.5 sec window hop) and training a classifier per time window produced a curve of accuracy vs. time window indicating how well the outcome was decoded at the population level above chance (Δaccuracy) in a specific session (Figure 3D and Figure S2B). For the control group, the population activity became predictive gradually from second session and onward at the post movement time window (approximately 5-6 seconds after the go cue), where for the manipulated group outcome was successfully decoded in later sessions (6^th^-7^th^), after CNO was lifted (Figures 3D and Figure S2B).

We next addressed the question whether the population encoding of outcome gradually transforms towards a fixed configuration, or rather the population representing outcome varies across training sessions being specific per training session. Toward this end, we used the models trained for the expert animals (7^th^ sessions) at the last analysis window (go cue +7.5 sec) and applied them to predict outcome signals in the neuronal activity of previous training sessions. The Δaccruacy (Figure 3E) as a function of training session showed that expert models were unable to decode outcome from ensemble activity of naïve animals (1^st^ training session). However, with training, the accuracy of the expert model in decoding outcome increased, and became significantly above chance from the 4^th^ session and onward (p=8.5·10^-4^, p=1.0·10^-4^, p=6.3·10^-5^, p=4.0·10^-9^ for sessions 4-7 respectively, tailed t-test, n=12). This monotonic increase in accuracy confirmed our hypothesis that encoding of outcome by population activity progresses towards a specific profile with training. For the manipulated group the accuracy of the expert model begun to rise only when CNO is lifted and became significant at the 6^th^ session (p=0.013 for the 6^th^ session and p=7.4·10^-5^ for the 7^th^ session, tailed t-test, n=7).

Collectively these results show that outcome representation by layer 2-3 PNs network is not inherent (as it does not exist in naïve animals) but acquired and gradually transforms towards a population activation profile through training, and that dopaminergic VTA projections to M1 play a fundamental role in facilitating this outcome network learning process.

### Functional network connectivity reorganization of layer 2-3 PN during motor learning

In addition to the changes in activity dynamics and population outcome representation throughout the training process, we set out to investigate the changes in the functional network connectivity configuration that evolves during the motor learning process. To get insight on the changes in the functional connectivity within the layer 2-3 network during the training sessions, we extracted a correlation matrix per trial for every training session based on pairwise Pearson’s correlation coefficients between all possible pairs of ROIs (Figure 4A). We next calculated the centroid of all trial-based correlation matrices of a given session, creating a single matrix for the different trials of a session. As correlation matrices are Semi-Positive Definite, they are located on a curved manifold and do not obey Euclidean geometry. Therefore, instead of using a Euclidean average, we used Riemannian geometry to extract their centroid on the curved manifold^49,50^ (see Methods). Overall, this analysis produced a single matrix per training session, the Riemannian centroid, representing the average correlational structure of the network in a given training session (Figure 4A).

**Figure 4.**
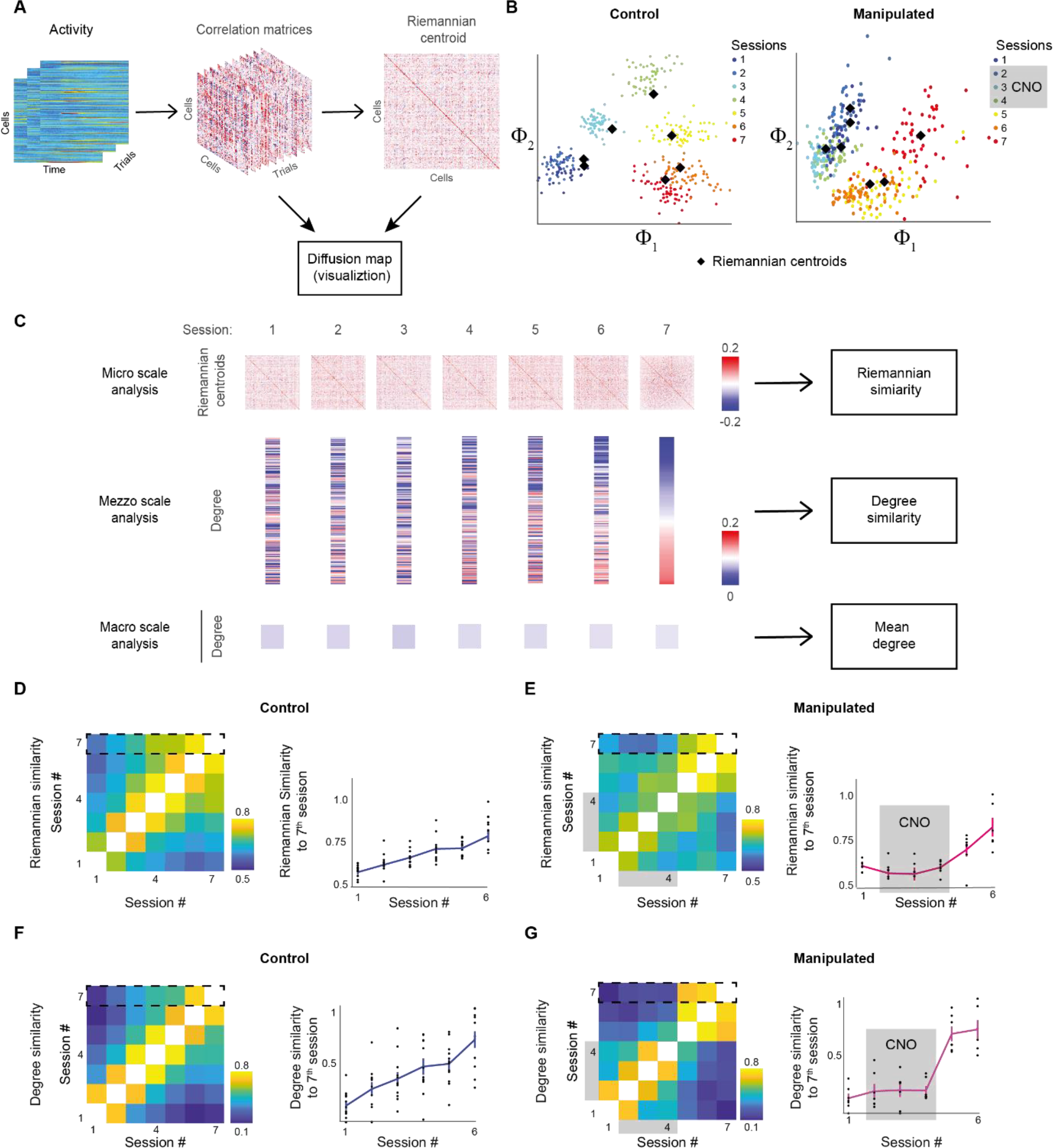
Transformation of the functional connectivity of layer 2-3 network during motor learning and the role of VTA dopaminergic projections to M1 in the process. **A**. Schematic illustration of the analysis pipeline. Construction of correlation matrices of the activity between pairs of neurons per trial, used to evaluate the Riemannian centroid representing the functional connectivity of the network in a given training session. **B**. Example of a two-dimensional diffusion embedding of trial-based correlation matrices (each dot represents a trial and colors indicate the training sessions) and their Riemannian centroids (black diamond) for a control (left) mouse and a manipulated mouse (right, CNO sessions 2, 3, 4). **C**. A schematic description of subsequent analysis in three levels: Micro, Riemannian similarity accounting for pairwise relations between all ROI pairs; Mezzo, degree calculation expressing the strength of connectivity of each cell to all other cells; Macro, mean degree expressing the global connectivity of the network. **D**. Riemannian similarity extracted from centroid matrices across different training sessions. Left, heat maps indicate similarity between pairwise sessions, averaged across animals. Right, Riemannian similarity (mean±SEM) comparing all training sessions to expert (7^th^ session), for the control group. **E**. Same as D for the manipulated group. **F**. Same as D for degree vectors through training sessions for control group. **G**. Same as F for the manipulated group. For D-G, black dots are population data (per animal), blue and pink are mean (±SEM) across animals, n=12 for controls and n=7 for manipulated mice.

For illustration purposes we first computed a low-dimensional representation for the trial-based correlation matrices and their centroids for a control animal and a manipulated animal using diffusion maps^51,52^ combined with Riemannian geometry (Figure 4B, see Methods), ultimately obtaining a visual representation of each session, where each trial was represented by a colored dot and the whole session by its Riemannian centroid (black)^37,38^. In the control animal, the embedded correlations were mainly grouped by the training session, that is the trials within each training session had similar connectivity matrices (up to some intrinsic trial variability), whereas trials from different sessions had different connectivity matrices. Furthermore, with training we observed a typical trajectory of transformation of the connectivity matrices indicating an evolution of the layer 2-3 neurons’ functional connectivity that occurs along the training process. For the manipulated group, the embedded correlation matrices did not differentiate well between the trials of the training sessions with CNO (2^nd^-4^th^) which were grouped together with the 1^st^ session and only in the following (5^th^-7^th^) sessions a differentiation developed. These results indicate the essential role of M1 dopaminergic projections in changing the functional connectivity profile of the network during learning.

We quantified the changes of the functional network connectivity with training, at three levels of analysis (Figure 4C). First, the detailed micro level of individual functional connections, where we took into account the specific identity of pairwise neuronal connectivity. Second the mezzo level of individual neurons, where we computed the degree of connectivity of each neuron with other neurons in the network regardless of the identity of the connection, and thereby quantifying the strength each neuron is connected to the network. Third, the macro level of the entire network, where we evaluated the average functional connectivity of all neurons in the network. For these three levels of analyses, we evaluated the functional connectivity of the network in each session as the Riemannian centroid matrix of all related trial-based correlation matrices (Figure 4A, see Methods).

To capture the transformation in functional network connectivity at the micro individual connection level, and to account for pairwise relationships between cells, we evaluated the Riemannian similarity between the Riemannian centroid matrices (Figure 4A) in different training sessions (Figures 4D-E, see Methods). In the control group, the Riemannian centroid connectivity matrices were most similar to subsequent training sessions, where similarity to the 7^th^ session was gradually rising (R^2^= 0.55, p=1.2·10^-13^, linear fit), while for the manipulated group, this process was observed only after CNO was lifted during the training sessions. The analysis at the micro level demonstrated that the functional connectivity reorganizes during the training and that the specific pairwise connectivity footprint each neuron makes in the network is important and evolves with training. Moreover, our finding shows that this reorganization process required dopaminergic activation in M1.

Next, we investigated whether the reorganization of the individual functional connections also impacts the average functional strength each neuron is connected in the network. It is possible that individual connections will change but the average connectivity per neuron will remain constant due to a balance between strengthening and weakening of connections of the neuron. To test this, we applied the mezzo level analysis and computed the degree connectivity of each neuron at a given training session. The degree (overall connectivity) of a neuron was calculated by summing over the rows of each Riemannian centroid connectivity matrix, resulting with a vector expressing the extent each cell is correlated with all other cells within the network in a given training session (Figure 4C) and compared the values of degree across training sessions (Figures 4F-G; see Methods). We found that the degree profile of the network transformed toward the expert configuration (7^th^ training session) in the control group. No such transformation was observed for the manipulated group in training sessions 2-4 where CNO was applied, and was resumed once CNO was lifted during the consequent training sessions. This finding indicates that the reorganization of the network during learning also involves changes in the overall strength of connectivity of individual neurons to the network and verified that dopaminergic inputs at M1 were essential for this transformation.

Finally, at the macro level, we tested whether the global connectivity of the network changes during the training sessions. We compared the average degree across all cells per training session (Figure 4C) and found that the mean degree of the network did not exhibit a significant change with training (p=1, n=12, for control group, p=1, n=7 for the manipulated group, one way ANOVA test). This result indicates that despite the reorganization of functional connectivity at the micro and mezzo levels, the overall connectivity level of the network remained unchanged. This result may be explained by the fact that some cells increased, and some decreased their connectivity such that the overall connectivity in the network did not change with training.

Taken together, our results indicate that layer 2-3 network undergoes a reorganization towards a specific functional connectivity configuration typical to the expert with training, which critically depends on activation of the VTA dopaminergic projections to M1. The expert configuration depends on both the specific connectivity profile of each neuron and on the strength of connectivity of each neuron within the network.

## Discussion

In this study, we examined the plasticity changes in the layer 2-3 PNs network of M1 during the learning process of a dexterous motor task, and the role that direct dopaminergic inputs to M1 play in this process. We found that while the average activity of layer 2-3 PNs did not change significantly during the training process, the activity kinetics along the trial changed and became more similar to the “expert” configuration of the 7^th^ day of training. Moreover, while we did not observe changes in the average functional connectivity of layer 2-3 PNs, we observed a gradual reorganization of the specific functional connectivity profile of neurons and the strength each neuron is connected in the network, which became more similar to the final “expert” functional connectivity configuration. The fact the average connectivity of the network remains stable probably reflects homeostatic mechanisms that maintain the overall excitability of the network. The monotonic transformation of activity kinetics as well as network functional connectivity during learning were critically dependent on the VTA dopaminergic inputs at M1, as silencing these projections during the learning sessions blocked both motor learning and plasticity changes in the reorganization of activity and functional connectivity of layer 2-3 PNs network. In addition, we found dopaminergic VTA projections to M1 are crucial in encoding outcome of the learnt task at the single cell and population of layer 2-3 neurons. This VTA dependent reorganization of specific functional connections during learning can serve to generate specific subnetworks (ensembles) that underlie memory.

### Layer 2-3 network organization in M1 during motor learning

We report that the averaged total layer 2-3 network activity did not change significantly during learning of the hand reach task. This is consistent with a previous study, that showed stability in the fraction of active layer 2-3 neurons during a lever press task^53^. However, despite this seemingly stable network activity a deeper look revealed learning was associated with major reorganization of both the neuronal activity dynamics and the network connectivity. During learning, the activity dynamics of neurons along the trial changed and transformed to the expert activity configuration adopting a typical three phasic kinetics^13^.

In addition to activity dynamics, we examined the effect of learning on functional connectivity. During most experimental paradigms, the access to real connectivity levels of neurons in networks is extremely limited. Thus, a common practice is to evaluate the functional connectivity by measurements of pairwise correlations between neuronal signals for various signals such as functional magnetic resonance imaging (fMRI)^54,55^, calcium imaging or single unit recordings^56^ where configurational structure of the network is compared across time or across subjects through centrality measures such as degree^57–61^.

In this work we use a similar practice, where we evaluate the functional connectivity between individual neurons using the correlation matrix, calculated per trial. Here we developed a geometric approach pipeline for tracking changes in correlational structure throughout the training process. In this new approach we calculate the Riemannian centroid across trials, to express the correlational structure of all neuronal pairs, in a given session. We found that while there were no significant changes in the overall connectivity of the network (globally averaged degree centrality), the learning process induced a significant change in the functional connectivity of the individual neurons and their total connections in the network thus keeping a homeostatic level of connectivity while introducing specific task related connectivity changes. The correlational structure expressed by all pairwise relations gradually transform and converges towards an “expert” configuration indicating that the learning influences specific connections in the network.

Taken together our findings support four processes that occur during learning. First, formation of new specific connections between neurons and thereby probably forming specific ensembles for storing the new task^62,63^. Second, to maintain the overall excitability of the network, weakening of other unrelated functional connections occurs. Third, shaping of a new tri-phasic dynamic of the neuronal response^13^. This is probably mediated both by the formation of new connections between excitatory neurons, and probably by connectivity changes with inhibitory interneurons^64^. The activity dynamics and connectivity of interneurons were not examined in this study, yet it is important to stress that interneurons in M1 are being innervated by the VTA^65^ possibly contributing to the tri-phasic dynamics of the layer 2-3 activity. Finally, learning was associated with the development of a specialized subpopulation of neurons that encoded task outcome. We hypothesize that these outcome signals are crucial for reinforcement motor learning^13^, providing feed-back for future adaptations.

Our results are in line with the increased temporal correlations of pairs of neurons reported for anterolateral (ALM) and posterior medial motor cortexes during learning of lick no-lick odor task, though correlations were generally low, and the network correlative configuration was not studied^24^. In addition, our results are also consistent with the previously reported increased stability of the temporal activity pattern of neurons and the relationship between activity and movement in pairs of trials that became more consistent with learning of a lever press task^22,53^.

Our results, however, are inconsistent with two previously described findings with regard to layer 2-3 neurons during motor learning. First, a previous study that reported unchanging total predictive decoding accuracy of the motor behavior by both the neuronal population or single layer 2-3 neurons of M1 during lever pull motor learning^44^, a property that was attributed specifically to layer 5a but not 2/3^44^. Second, our results are also inconsistent with the notion that the single neurons are unstable in their activity during trials, tuning or predictive decoding of behavior during the learning sessions^22,24,44,53^. We show that increasing fraction of neurons gradually change and converge toward the “expert” activity configuration where each neuron adopted gradually its specific activity pattern during the learning process, which can potentially explain the gradual transformation towards the expert configuration seen at the network functional connectivity level.

### The role of M1 VTA projections in motor learning: behavioral and layer 2-3 activity

Previous work demonstrated that the main source of dopaminergic projections to M1 originate from the meso-cortical dopaminergic system, the VTA. Further, this pathway was shown to be necessary for hand reach motor skill learning^29^ as eliminating the dopaminergic terminals with 6-OHDA in M1 or blocking D1 and D2 receptors affected the motor skill learning^66,67^. Consistent with previous works, behaviorally, we found that dopaminergic inputs to M1 are essential for motor learning as silencing these inputs locally in M1 prevented learning progression^29,34,68^. However, in contrast to a previous study that showed VTA lesioning did not affect learning within session rather only between sessions^29^, we show that inhibiting VTA terminals with DREADDs in M1 halted both within and between sessions motor learning, preventing mice from increasing the success rate in the CNO sessions. In addition, our analysis of the motor behavior during the hand reach task revealed that inhibition of the direct dopaminergic projection also decreased the number of task related movements and delay time from go cue to first lift, indicating the importance of direct dopaminergic projections on learning not only through success rates but also on fine behavioral parameters. These findings are in line with recent work showing a role of dopamine in coordinating fine reaching skilled movements^69^ though in this case midbrain dopaminergic neurons in the substantia nigra pars compacta (SNc) and not VTA dopaminergic neurons were manipulated.

Interestingly, previous studies have reported the importance of dopaminergic SNc afferents to the striatum in motor learning^70–73^. Thus, motor learning probably requires concerted activation of both direct dopaminergic signaling to M1 via the VTA and dopaminergic signaling to the striatum. It is yet unclear whether the two different pathways carry different information and/or are responsible for different aspects of learning or whether learning requires a double gate dopaminergic mechanism. The fact M1 receives both a general reward prediction error signals^1,27^ and develops a specific task related outcome signal as part of the learning process may be related to this question. Further studies are required to unravel this interesting and important question.

While the role of direct dopaminergic VTA afferents to M1 on learning was described previously at the behavioral level, information regarding the role of dopaminergic VTA projections to functional connectivity and activity dynamics in M1 during learning was lacking. In this study we attempted to close this gap and show that for both single neuron dynamics and network functional connectivity reorganization to emerge, and for the concomitant motor learning to proceed, activation of direct VTA dopaminergic inputs to M1 is essential.

It is interesting to note that the dopaminergic VTA axons project to a wide range of areas in the brain^74^, with the majority of projections innervating subcortical regions, primarily the nucleus accumbens (medial and lateral) and amygdala^75–78^. VTA is also known to innervate other cortical regions, especially the prefrontal cortex^75^, however, other cortical regions including M1 are only scarcely innervated. Here we show that despite this scarce innervation, dopaminergic VTA projections to M1 are vital for motor learning and for the reorganization of the M1 network during the training process.

Dopamine is thought to play a major role in reinforcement learning providing a reward prediction error signal^79^, comparing between expected and experienced outcome which than leads to strengthening of the appropriate synapses via plasticity mechanisms^80,81^. In this study we did not address the cellular mechanisms by which dopamine participates in reorganization of the M1 network during learning. Previous studies showed that the dopaminergic innervation to M1 from the VTA is important for inducing LTP in M1 layer 2-3 synapses^34^ in line with the findings that Horizontal connections of layer 2-3 motor cortex undergo LTP during forelimb motor skill learning^26,68^. At the structural plasticity level task related spine appearance and elimination in layer 2-3 neurons during motor learning was shown^23,25,53^. Moreover, in layer 5 neurons spine turnover was shown to depend on direct mesocortical projections to M1^82^. It is likely that one or more of these mechanisms are responsible for the effects we describe in this study. Thus, our findings together support the notion that the reorganization at the single cell and network levels of M1 are supported by dopamine released during the motor learning process from the VTA.

## Acknowledgments

We thank Y. Schiller for helpful discussion and for comments on the manuscript. We also thank B. Engelhard, Y. Gutfreund and O. Barak for helpful comments on the manuscript. Funding: This study was partially supported by the Israeli Science Foundation (J.S.), Prince Funds (J.S.) and Rappaport Foundation (J.S.).

## Author contribution

Conceptualization, J.S.; Methodology J.S., A.G., H.B., R.T.; Software, H.B., A.CR.; Formal Analysis, A.G., J.S., H.B., A.CR., R.T., S.S.; Data Curation, H.B., J.S.; Writing, J.S., A.G., H.B., R.T.; Visualization, H.B.,

A.G., A.CR.; Funding Acquisition, J.S.

## STAR Methods

### Key resources table

| REAGENT or RESOURCE | SOURCE | IDENTIFIER |
| --- | --- | --- |
| Bacterial and virus strains |  |  |
| AAV.CamKII.GCaMP6s.WPRE.SV40 | James M. Wilson lab (unpublished) | Addgene, viral prep #107790-AAV9 |
| pAAV-hSyn-DIO-hM4D(Gi)-mCherry | Krashes et al., 2011 | Addgene viral prep #44362-AAV2 |
| Experimental models: Organisms/strains |  |  |
| Common Name: DAT-cre | Zhuang X et al., 2005 | JAX stock #020080 |
| Software and algorithms |  |  |
| MATLAB | MathWorks | Version 2019a, Statistics and Machine Learning Toolbox, https://www.mathworks.com/ |
| LabView | National Instruments | Version 2018, https://www.ni.com/en-il/shop/labview.html |
| JAABA | Kabra et al., 2013 | http://jaaba.sourceforge.net/ |
| LabView interface homemade code | Schiller's Lab, Technion, Israel | https://bitbucket.org/natecermak/mouse-task-controller/src/master/TwoPhotonTrialController_reachTask.vi |
| Classification, dynamical and statistical analysis | Schiller's Lab, Technion, Israel | https://github.com/hadasbenisty/networks_learning_M1 |

## RESOURCE AVAILABILITY

### Lead Contact

Further information and requests for resources and reagents should be directed to and will be fulfilled by the Lead Contact, Jackie Schiller.

### Materials Availability

This study did not generate new unique reagents.

### Data and Code Availability

The datasets supporting the current study have not been deposited in a public repository because they are large files, but are available from the corresponding author on request. The code generated in this study is available at GitHub.

### Experimental model and subject details

All animal procedures were performed in accordance with guidelines established by the NIH on the care and use of animals in research, as confirmed by the Technion Research Campus Institutional Animal Care and Use Committee. Adult male Slc6a3 (DAT-Cre) mice were used in this study. Animals were housed in a 12:12 hours reverse light: dark cycle. For behavioral training and experiments, food intake was limited to 2.65–3 g/day with ad libitum water.

### Experimental design

To investigate the activity and network dynamics in M1 during motor learning, we trained mice to perform a head-fixed version of the hand reach grasping motor task. After the first successful hand reach to pellet attempt, all subsequent training sessions (1-7 sessions) were performed with simultaneous two-photon calcium imaging (control group). Chronic calcium imaging was performed with the genetically encoded calcium indicator GCaMP6s, which was introduced to the cells via a viral vector injection. The resulting permanent expression of the indicator allowed us to monitor the activity for many weeks. We selectively labeled layer 2-3 PN using a specific promotor (CaMKII). We implanted a chronic window over the M1 forelimb area that allowed us to image the activity for long time periods. For silencing the VTA activity we used chemogenetic inhibition using DREADDs which was introduced to the cells via viral vector injection (manipulated group). We selectively labeled dopaminergic neurons in the VTA using Cre-dependent viral expression in DAT-Cre mice. The manipulation was done for three consecutive training sessions (2, 3 and 4) by injecting CNO locally to M1 via an access port^83^ which allowed specific inhibition of the dopaminergic projections from VTA to M1.

### Viral injections and cranial window surgery

The procedures were performed on 2-3 months old mice, under isofloren anesthesia (4% for induction and 1.5-2% for maintenance during the surgery). Mice were secured on a stereotaxic apparatus (KOPF), and a heat pad was used to maintain body temperature at 36-37 °C. Pre-operation medications were administrated subcutaneously for analgesia, Ketoprofen (5 mg/Kg) buprenorphine (0.1 mg/Kg). The scalp was shaved and cleaned with ethanol. The skull surface was exposed after a subcutaneous injection of 2% Lidocaine. Viral injection was performed using a hydraulic manipulator (M0-10 Narishige) via thinned skull at two points: 1. M1 forelimb area^10^ (0.6 mm anterior and 1.6 mm lateral to Bregma) at three depths (150 μm, 250 μm, 350 μm, with 70 nl each depth). For calcium imaging at M1, we injected AAV.CamKII.Gcamp6s. 2. VTA bilaterally (3.1 mm posterior, 0.3 mm lateral to Bregma; depth of 4200 μm, volume of 120 nl for each side). For VTA inhibition we used DREADDs AAV2-hSyn-DIO-hM4D(Gi)-mCherry. After injections we performed a circular craniotomy over the M1 forelimb area (0.6 mm anterior and 1.6 mm lateral to Bregma) and implanted a 3mm diameter laser-cut glass optical window with a 1mm access hole which we sealed with biocompatible silicone for future local injections. The window was sealed with superglue and dental cement. Next, a custom-made 3D printed plastic headpost^13^ was affixed to the skull with dental cement. Mice were injected with ketoprophen (5 mg/kg) and buprenorphine (0.1 mg/kg) administered subcutaneously for 2-days post operation and were left to recover from the surgery for two weeks with *ad libitum* food and water.

### Local CNO injections

For the manipulated group, we injected the CNO locally in M1 immediately before starting the training sessions. The injections were done via an access port in the cranial window which was sealed with biocompatible silicone^83^ using hydraulic micromanipulator with a glass pipette that was inserted into the cortex with an angle to reach the specific imaged area in M1. Approximately 500 nl of CNO (100 μM) was injected. The experiments started 5 min after the injection of the CNO and lasted for 30-40 min.

### Behavioral training

After recovering from the surgery mice were restrained to 2.65-3 gr/day and *ad libitum* water. Training started when mice reached 80-90% of their initial body weight. Mice were habituated to head fixation in a custom-built apparatus^84^ in dark and quiet conditions, monitored by a webcam. We trained to reach for a food pellet (20 mg; Test Diet; St Louis, MO) from a rotating table placed in front of the animal and driven by either a NI USB DAQ device or a Teensy microcontroller driven by custom-made LabVIEW software. An auditory tone (200 ms, 1 kHz) was used as a cue during plate rotation. Initially, the rotating table was placed directly below the mouth allowing the animal to tongue reach the pellet upon the tone. We gradually placed the table in further position until the first successful hand reach. Thereafter, training was combined with two-photon calcium imaging in 7 subsequent sessions. The first training day with imaging was called “session 1”.

### Two-photon calcium imaging

Images (512 X 512 pixels) were acquired using a two-photon microscope (Bruker) equipped an 8 kHz bidirectional resonant galvo scanner and a Nikon 16X CFI Plan Fluorite objective (NA 0.8, 3 mm working distance), controlled by the software package PrairieView 5.3. GCaMP6s was excited at 940 nm using a femtosecond pulsed laser (InSight X3, Spectraphysics). Emission light was detected by a GaAsP photomultiplier tube (Hamamatsu).

Each trial was 12 sec with an auditory cue that was presented in the 4^th^ second. The same field of view was imaged over all sessions in the same animal. Behavioral performance was monitored at 200 Hz using two cameras (side and front view; Flea3 FL3-U3-13Y3M, PointGrey). The timing of two-photon calcium imaging, behavioral task, and video recording were coordinated via a National Instruments board (PCI-6110), using custom-made software written in MATLAB.

### Histology

At the end of experiments, mice were deeply anesthetized and transcardially perfusion with PBS 0.1% followed by paraformaldehyde 4%. The brains were removed and put in fixative solution (paraformaldehyde 4%) for 48 hours, and later stored in PBS 1% solution. Coronal sections (100 μm thickness) were made. The sections were mounted on slides embedded with Fluoroshield containing DAPI (Sigma) and images using Olympus stereoscope (MVX10M).

## QUANTITATION AND STATISTICAL ANALYSIS

### Behavioral data analysis

We labeled the behavioral events (Lift, Reach, Grab, Supinate, At-Mouth using a semi-supervised software^45^ as follows: Lift – a vertical movement of a forelimb from the perch or the table, lasting till peak height is reached; Reach – a horizontal movement towards the pellet; Grab – starts when animals spread their fingers, ending when fingers are wrapped into a fist. Supinate – the forepaw is rotated outward while being brought towards the mouth; At-Mouth – fist is close to the mouth.

To characterize behavior throughout training we segmented the data into sequences of behavioral events and counted the number of inter-sequence transitions, per trial.

### Two-photon data analysis

The fluorescence data acquired by the two-photon microscope was first registered to correct for brain motion artifacts. Our registration method was based on^85^, using Fourier transform based correlation between two successive images. The maximal value position in the correlation image specifies the relative shift between the two images; we designate them *u*_*t*_ and *v*_*t*_. This method requires a template specification and matching against an image stack. The template image *I*_*temp*_(*x, y*) was defined as the average of all images in the selected trial over time.

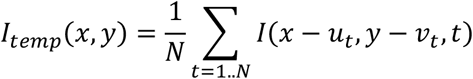

The set {*u*_*t*_, *v*_*t*_}, *t* = 1. . *N* is an image shifted in the XY plane after alignment. We initially start with *u*_*t*_ = 0, *v*_*t*_ = 0 and then update their values according to the registration maxima. This procedure is repeated several times, when each time we compute the new template *I*_*temp*_(*x, y*) using previously computed {*u*_*t*_, *v*_*t*_} for each image. Typically, this procedure converges after several iterations, in our case three iterations.

To align the imaging data over many trials, we used a similar technique, utilizing the previously computed averaged templates for each trial. For each trial *k*, we performed a single trial registration using the template algorithm for three iterations. To align the image data over many trials, we treated the final templates 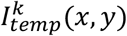 from each trial *k* as unaligned image data and repeated the same registration procedure to find offsets {*u*_*k*_, *v*_*k*_} for each trial. These offsets, along with previously found offsets {*u*_*t*_, *v*_*t*_}, account for the final image shift in the XY plane.

Regions of interest (ROIs) were detected manually using average fluorescence images and *ΔF⁄F* projection images, which highlighted active neurons. The pixels within each ROI were averaged for every frame. The ROI ‘‘mask’’ was used to detect the same neurons on multiple imaging sessions on different days.

*ΔF⁄F* was computed using the following formula:

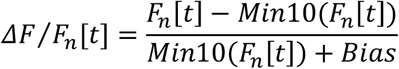

*Min*10(*F*_*n*_[*t*]) is a mean value of the lowest consecutive 10% values of the fluorescence signal *F*_*n*_[*t*]. This minimal fluorescence value was calculated both per trial and across the entire experimental session, with no significant differences in the results with these two calculation variants. A small bias factor in the denominator prevented zeros when the cell was completely silent.

### Statistical Analysis

In each experiment, the mouse repeated the hand reach task several dozen times. The data were analyzed using custom Matlab code, except for training and testing the SVM classifiers, for which we used the standard LIBSVM software^86^.

The imaging data collected at each session *l* are stored in a 3-dimensional matrix (tensor) ***X***^??^ of size *N*_*r*_ × *t* × ***T***^*l*^, where *N*_*r*_ is the number of neurons, *t* is the number of time samples per trial, and ***T***^*l*^ is the number of trials. *X*^*l*^(*i, j, k*) is the neuronal activity of the *i*-th neuron at the *j*-th time sample in the *k*-th trial related to the *l*-th session, where *l* = 1, …,7.

#### Activity of Cells Through Training

The overall average activity of the entire population of cells per training session was obtain as:

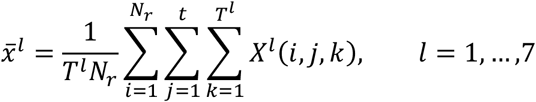

The average dynamics of cells in each session was obtained by averaging over trials as follows,

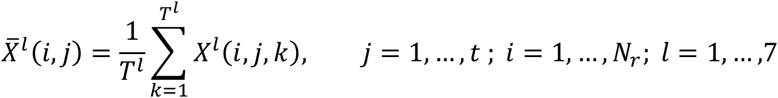

We obtained the correlation coefficient between the average dynamics of each cell in the 7^th^ session with its own dynamics in all other sessions:

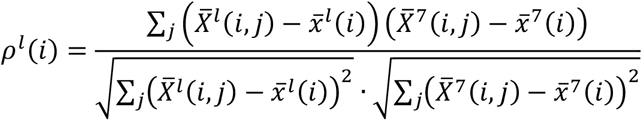

where 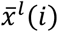 is the activity of the *i*-th neuron in the *l* session, averaged across time and trials.

#### SVM Classification of Ensemble Activity

To further quantify the activity profile of the cellular ensemble throughout training, we trained a linear SVM classifier^47^ to separate neuronal activity in trials related to the first session from the activity in trials related to the 7^th^ session. We used a sliding window of 1 sec with 0.5 sec hop. In every time window, we evaluated the average activity of each neuron at each trial,

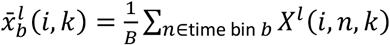

where *b* is the index of time window and *B* is the window length measured in samples. The activity of the network in each time window for each trial was represented by the following *N*_*r*_ × 1 vector:

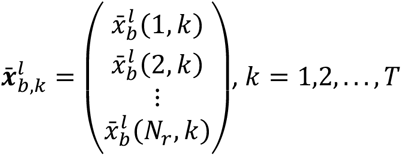

For each time bin, we extracted the set 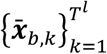, representing the averaged activity of the ensemble in all trials related to all 7 training sessions. We paired the activity of each trial with a label indicating whether it is associated with expert activity (7^th^ session) or with naïve activity (1^st^, session):

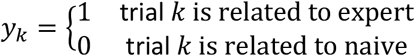

We used the standard LIBSVM toolkit^86^ to train a linear classifier to separate expert from naïve, per time bin, using 10-fold cross validation. We then applied the trained classifiers to estimate trials from sessions 2-6. For the 1^st^ and 7^th^ sessions, we used the output of the cross-validation procedure. For each session we evaluated the fraction of trials identified as expert, per time window *b*,

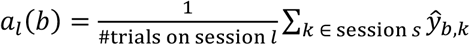

where 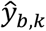 is the estimated label. Figure 2F presents mean±SEM values of *a*_*l*_(*b*), averaged across animals, for a time window centered at tone+1sec. Figure 2G presents the average values as a function of the time bin *b*.

#### PCA Trajectories of Network Dynamics

We explore the evolution of the network in encoding outcome using Principal Component Analysis (PCA). We reshaped the imaging data tensor of a given session, *X*^*l*^, into a 2-dimensional matrix of *N*_*r*_ × [*t****T***^*l*^], where each row consists of the neuronal activity of a specific cell across all time samples and trials:

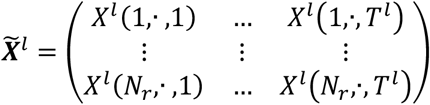

We computed the sample covariance of 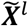 and extracted its eigenvalue decomposition to obtain *d* principal components explaining 95% of the variance of 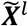. Finally, we reshaped the new representation into a *d* × *t* × ***T***^*l*^ tensor of the principal components of *X*^*l*^. Figure 3B presents the first 3 principal components, averaged according to outcome, computed per session for two animals.

#### Significance in Separation

Let ***S***_*suc*_ be a *d* × *t* × ***T***^*suc*^ tensor of all trajectories related to successful trials in each session, and ***S***_*fail*_, a *d* × *t* × ***T***^*fail*^ be the corresponding tensor for failed trials. First, we computed the average trajectories for success and for failure, *μ*_*suc*_(*n*) and *μ*_*fail*_(*n*), and the corresponding sample covariance matrices, *Σ*_*suc*_(*n*), *Σ*_*fail*_ (*n*):

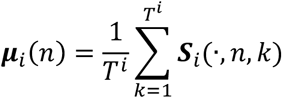

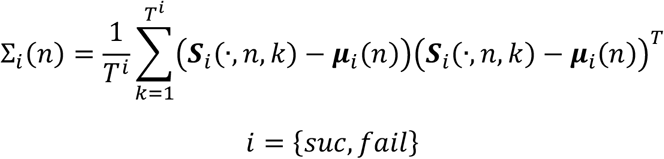

Second, we calculated the separation between success and failure dynamics using:

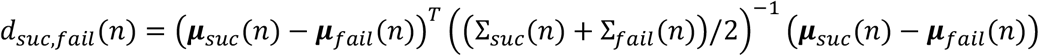

Third, we shuffled the trial outcome labels and then re-computed the distances. We repeated this computation 1000 times per session and obtained the empirical probability for the randomized distance to be larger than the actual distance, denoted by 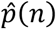. Finally, for each training session *l*, and time point *n*, we counted how many animals yielded a significant difference, 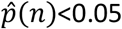, as presented in Figure 3C.

#### Classification of Outcome Per Training Session

We further quantified outcome separation emerging with training based on the predictive power of the imaging data tensor *X*^*l*^ using a linear SVM classifier^87^. The classification accuracy was evaluated for each training session, using a sliding window of 1 second with 0.5 second hop. In every time window *b*, we evaluated the average activity of each neuron at each trial, 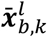, as defined above. We then trained a binary SVM classifier (linear) to separate successful trials from failed trials, per time window, using the standard LIBSVM toolkit^86^. Training and testing were performed using a 10-fold cross-validation procedure. As success rate varies with training and across animals, we evaluated the decoding power of the trained classifiers as the difference between accuracy and chance level, calculated per training session.

#### Classification of Outcome Using Expert Model

We applied the SVM models trained to separate success from failure based on the activity of expert animals, at the last time window (go cue+7seconds) to data extracted from previous sessions (last time window as well).

#### Riemannian Centroid of Correlation Matrices

We computed the pairwise Pearson’s correlation coefficients based on the activity of cells at each trial:

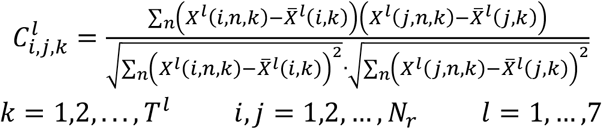

where 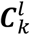 is the Pearson correlation matrix of the *k* −th trial in the *l*-th training session. The resulting Pearson correlation matrices 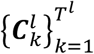 are Symmetric Positive Semi Definite (SPSD) matrices and therefore do not obey Euclidean geometry. To correctly evaluate the overall connectivity in a given session and compare connectivity across sessions, we utilized the particular geometry proposed in^37^ and^38^ as follows. Consider two Pearson correlation matrices *C*_*k*_ and *C*_*l*_ of rank *r*. We decomposed each of these matrices such that:

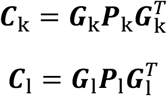

where ***G***_k_ and ***G***_1_ are *N*_*r*_ × *r* matrices with orthogonal columns that lie on Grassmann manifold^89^ and ***P***_k_ and ***P***_1_ are *r* × *r* Symmetric Positive Definite (SPD) matrices that lie on the SPD cone manifold^40^. The Riemannian SPSD distance proposed in^37^ is given by:

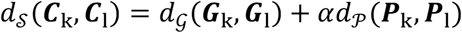

where α > 0 is a tunable hyperparameter, *d*_*𝒢*_ is the geodesic distance on the Grassmann manifold^89^, and *d*_𝒫_ is the geodesic distance on the SPD manifold^40^. The centroid of the Pearson correlation matrices in each training session was calculated as their Riemannian (Frechet) centroid^38^:

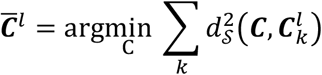

#### Diffusion Embedding of SPSD Matrices

We made use of this Riemannian distance to illustrate the development of the correlations of cells along the training sessions, by incorporating it into Diffusion Maps^52^, a non-linear dimensionality-reduction algorithm. Using Diffusion Maps, the collection of Pearson correlation matrices 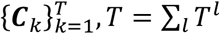 corresponding to trials from all experiments were embedded into a Euclidean space. First, we constructed a ***T*** × ***T*** affinity matrix, whose elements are given by

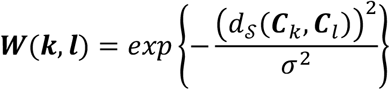

where σ is a scaling tunable parameter. This affinity matrix could be viewed as the weight matrix of the edges of graph, whose nodes are the trials. By normalizing the rows of ***W***, we obtained a row stochastic matrix ***P***,

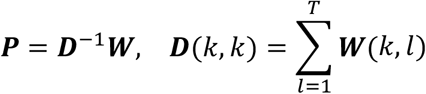

Then, we applied eigenvalue decomposition to ***P***,

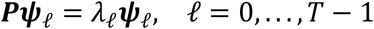

where *ψ*_*ℓ*_ and *λ*_*ℓ*_ are the eigenvectors and eigenvalues of ***P***. Finally, we defined *Ψ*_*k*_ as the diffusion map of the *k*th trial to a Euclidean space *R*^*d*^ by

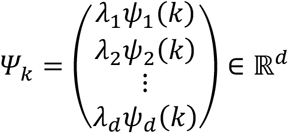

where *d ≤* ***T*** − 1. The Euclidean distance between trials in the embedded space constitutes the diffusion distance, which is related to a distance between the respective transition probabilities on the graph.

#### Riemannian Similarity

We quantified the distance between centroids related to different training sessions using the Riemannian SPSD distance, 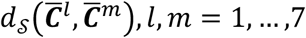 described above. As the number of cells greatly affect the dynamic range of these distances, we z-scored each matrix (per animal) to produce 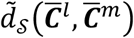. Finally, the similarity between centroids was obtained as:

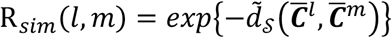

#### Degree Similarity

We obtained the degree of each neuron per training session *l*:

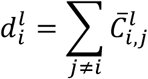

We calculated the distances between degree vectors ***d***^*l*^ related to different days as:

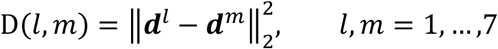

We then z-scored each matrix D per animal to obtain 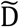 and extracted the degree similarity presented in Figure 4F-G as:

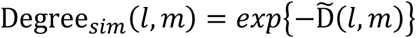

## Supplemental information

**Figure S1:**
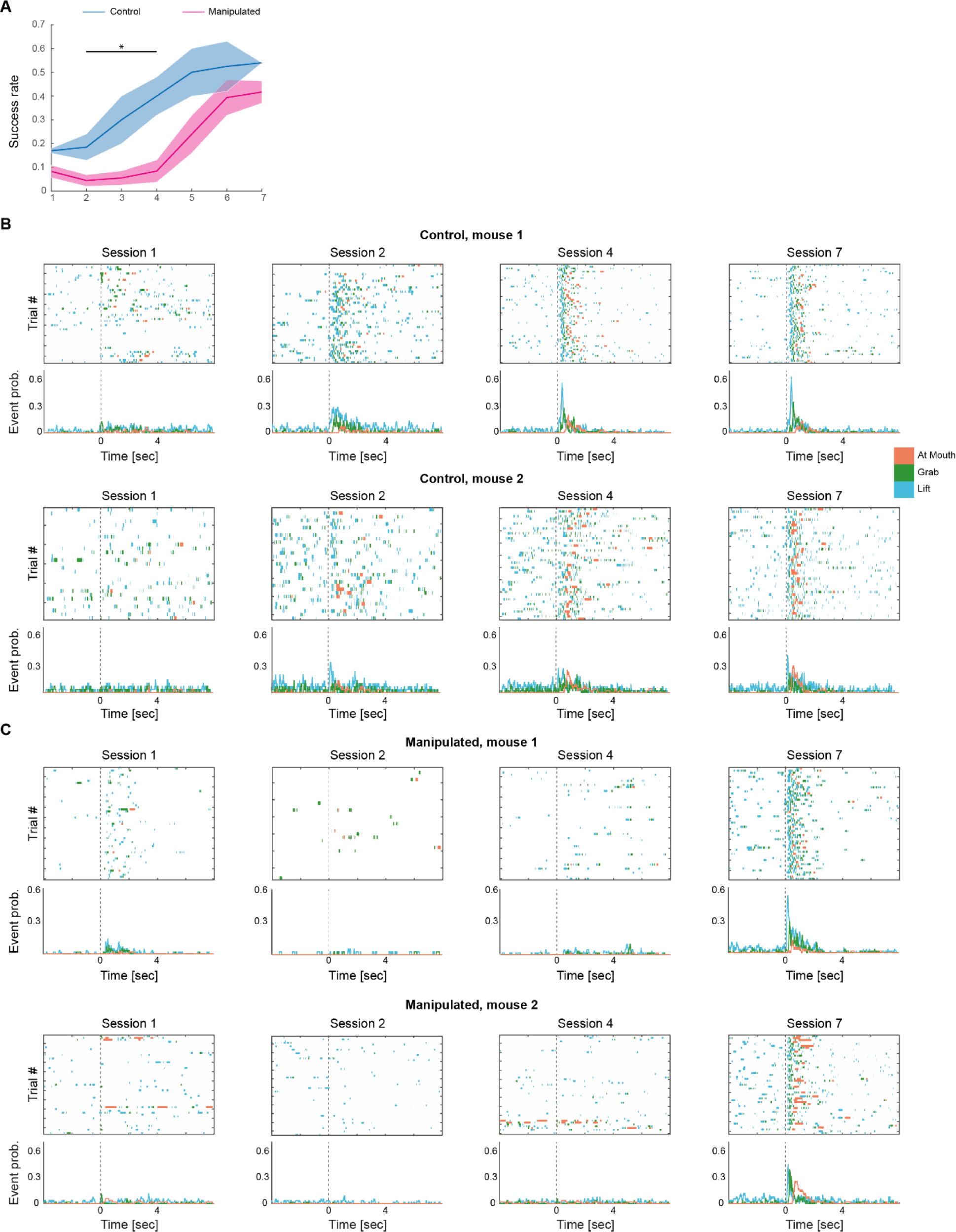
Control for the CNO injection, and example ethograms for control and manipulated mice. **A**. Success rate (mean±SEM) as a function of training sessions for the control group (n=2) injected with CNO at training sessions 2-4 (blue) and for the manipulated group (n=7) injected with CNO at training sessions 2-4 (pink). The injected control group demonstrates progression during the CNO session and reaches success rates that are significantly higher compared to the manipulated group (p=1.8·10^-4^, unpaired t-test for the three CNO injected training sessions between the viral injected and non-viral injected groups). **B**. Example ethograms for two control mice along four training sessions (1, 2, 4, 7-expert). Top, ethograms presenting ‘Lift’, ‘Grab’ and ‘At Mouth’ as a function of time (seconds) and trials on sessions 1, 2, 4 and 7. Bottom, empiric probabilities for events corresponding to ethograms. **C**. Same as in B but for two manipulated mice (CNO at training sessions 2-4). * Indicates p<0.05.

**Figure S2:**
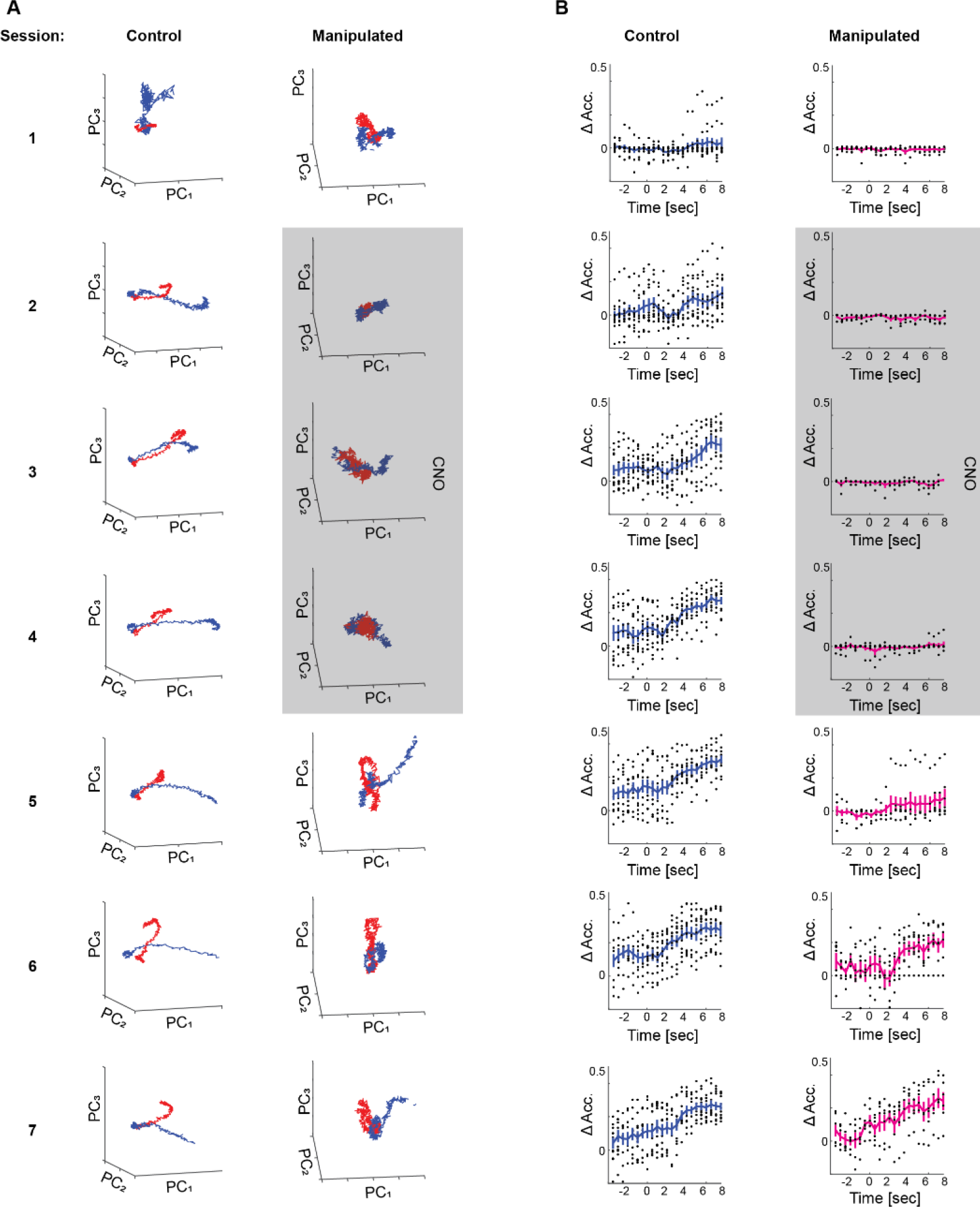
Network dynamics and population level success/failure separation. **A**. Example of temporal network dynamics during all sessions as captured by the first 3 principal components averaged across trials according to outcome: success (blue) and failure (red). Right column, control mouse; left column, manipulated mouse (grey box indicating CNO sessions). **B**. Population data showing delta accuracy rate (accuracy minus chance level) for classification of outcome based on the ensemble activity in a sliding window (window length 1sec, window hop 0.5sec), evaluated during all sessions. Right column, control group (12 mice); left column, manipulated group (7 mice). Black dots are population data (per animal), blue and pink are mean (±SEM) across animals. This figure demonstrates data from all training sessions (1-7), while in figures 3B, D data is represented for four selected sessions (1,2,4,7).

**Video S1: Motor performance during first session of a control animal**.

A video demonstrating the performance of the hand reach task during the first learning session of a control animal. This video is an example of a failure trial.

**Video S2: Motor performance during fourth session of a control animal**.

A video demonstrating the performance of the hand reach task during the fourth learning session of a same control animal as in Video S1. This video is an example of a successful trial upon two attempts.

**Video S3: Motor performance during seventh session of a control animal**.

A video demonstrating the performance of the hand reach task during the seventh learning session of same control animal as in Videos S1-2. This video is an example of a successful trial on first attempt.

**Video S4: Motor performance during first session of a manipulated animal**.

A video demonstrating the performance of the hand reach task during the first learning session of a manipulated animal. This video is an example of a failure trial.

**Video S5: Motor performance during fourth session of a manipulated animal**.

A video demonstrating the performance of the hand reach task during the fourth training session, i.e. third CNO injected session. Same manipulated animal as in Video S4. This video is an example of a failure trial.

**Video S6: Motor performance during seventh session of a manipulated animal**.

A video demonstrating the performance of the hand reach task during the seventh learning session of same manipulated animal as in Videos S4-5. This video is an example of a successful trial on first attempt.

